# Cyclic pentapeptide cRGDfK enhances the inhibitory effect of sunitinib on TGF-β1-induced epithelial-to-mesenchymal transition in human non-small cell lung cancer cells

**DOI:** 10.1101/2020.04.27.063776

**Authors:** Kyeong-Yong Park, Jiyeon Kim

**Affiliations:** Department of Integrated Material’s Development, CHA Meditech Co., Ltd, Daejeon 34025, Republic of Korea; Department of Medical Laboratory Science, College of Health Science, Dankook University, Cheonan 31116, Republic of Korea

**Author notes:** Corresponding author: Jiyeon Kim, Ph. D.; Department of Medical Laboratory Science, College of Health Science, Dankook University; 119, Dandae-ro, Dongnam-gu, Cheonan 31116, Republic of Korea.

## Abstract

In human lung cancer progression, the EMT process is characterized by the transformation of cancer cells into invasive forms that migrate to other organs. Targeting to EMT-related molecules is emerging as a novel therapeutic approach for the prevention of lung cancer cell migration and invasion. Traf2- and Nck-interacting kinase (TNIK) has recently been considered as an anti-proliferative target molecule to regulate the Wnt signaling pathway in several types of cancer cells. In the present study, we evaluated the inhibitory effect of a tyrosine kinase inhibitor sunitinib and the integrin-α_V_β_3_ targeted cyclic peptide (cRGDfK) on EMT in human lung cancer cells. Sunitinib strongly inhibited the TGF-β1-activated EMT through suppression of Wnt signaling, Smad and non-Smad signaling pathways. In addition, the cRGDfK also inhibited the expression of TGFβ1-induced mesenchymal marker genes and proteins. The anti-EMT effect of sunitinib was enhanced when cRGDfK was treated together. When sunitinib was treated with cRGDfK, the mRNA and protein expression levels of mesenchymal markers were decreased compared to the treatment with sunitinib alone. Co-treatment of cRGDfK has shown the potential to improve the efficacy of anticancer agents in combination with therapeutic agents that may be toxic at high concentrations. These results provide new and improved therapies for treating and preventing EMT-related disorders, such as lung fibrosis and cancer metastasis, and relapse.

## Introduction

Epithelial-to-mesenchymal transition (EMT) is a process in which closely packed epithelial cells with polarity become more motile and invasive and turn into spindle-shaped mesenchymal cells. In general, EMT can be seen in the complex process of transformation that epithelial cells must undergo to acquire mesenchymal cell characteristics during embryogenesis, development, wound healing, and organ fibrosis [1,2]. Notably, EMT-induced mobility and invasion potential play an important role in cancer metastasis to other organs. Because the metastatic process is a major cause of death and poor prognosis in cancer patients, suppression of signaling pathways involved in the EMT process is emerging as a new therapeutic strategy in cancer.

Non-small cell lung cancer (NSCLC) accounts for approximately 80-85% of total lung cancer and has high mortality due to invasion and metastatic ability [3]. Despite the development of chemotherapy, radiotherapy, and surgery, the 5-year survival rate for lung cancer has not increased significantly. In lung cancer progression, transforming growth factor (TGF)-β is a major cytokine that induces invasion and metastasis through an EMT process [4-7]. In the induction of EMT by TGF-β, TGF-β phosphorylates TβR-I by binding to transmembrane Ser/Thr receptors TGF-β type I (TβR-I) and type II (TβR-II), subsequently phosphorylating the downstream mediators Smad2 and [8,9]. The phosphorylated Smad2/3 complex recruits Smad4, which translocates to the nucleus and binds to transcription factors, such as Snail/Slug and Twist, to activate the TGF-β-reactive genes [10-12]. TGF-β-mediated non-Smad signaling pathways, including the Wnt/β-catenin signaling pathway, are also involved in the EMT process in several types of cancer cells [13-15].

In the last decade, Traf2- and Nck-interacting kinase (TNIK) has been reported as a first-in-class cancer target molecule [16]. TNIK kinase activity and expression have been shown to be involved in the maintenance of cancer growth in colorectal cancer, blood cancer, and lung adenocarcinoma [17-20]. Notably, we previously reported the potential of TNIK as an anti-cancer target molecule on the TGF-β induced EMT process, migration, and invasion of human lung adenocarcinoma cells [18]. In that study, TNIK inhibitor inhibited the TGF-β-induced EMT, migration, and invasion process through the suppression of Smad and non-Smad signaling pathways, including the TCF4/β-catenin involved Wnt signaling pathway. Although the association of TNIK with EMT in non-small cell lung adenocarcinoma has not been fully studied in many reports, inhibition of metastasis and invasion through TNIK inhibition is expected to play a role in increasing the therapeutic effect.

In this study, we evaluated the effect of sunitinib, a multi-targeted receptor tyrosine kinase inhibitor with high affinity for TNIK [21]. Sunitinib is a clinically approved small molecule drug that exerts anti-angiogenic and anti-proliferative activity against NSCLC through the inhibition of certain receptor tyrosine kinases, such as the platelet-derived growth factor receptors (PDGFRs), the vascular endothelial growth factor receptors (VEGFRs), and the stem-cell factor receptor KIT [22]. In addition, combination of sunitinib with an α_V_β_3_ integrin receptor-targeted cyclic-pentapeptide RGDfK (cRGDfK) enhances the anti-proliferative and anti-EMT effect in human lung adenocarcinoma cells.

The cRGDfK peptide has been identified as an integrin α_V_β_3_ inhibitor that acts as a targeting peptide for enhancing the permeability of tumor cells to drugs in anti-angiogenic cancer therapy [23]. The peptide ligand containing the arginin-glycin-aspartic acid (RGD) shows a strong binding affinity and selectivity to integrin α_V_β_3_ which sequence was discovered in fibronectin [24,25]. The RGD sequence is the cell attachment site with integrins that serves a recognition site for cell adhesion. The RGD-recognition sites were found in adhesive extra cellular matrix (ECM), blood, and cell surface proteins. Particularly, integrins recognize this RGD-containing sequence in their adhesion protein ligands [26]. Among the integrins, integrin α_V_β_3_ is consist of two subunits, α_V_ subunit and β_3_ subunit, which binds to RGD-containing ECM proteins such as vitronectin, fibronection and thrombopondin [27,28]. Based on these findings, linear or cyclic RGD-containing peptides have been developed as integrin α_V_β_3_ targeted ligands. Especially, cyclic peptide cRGDfK and cRGDyK showed high binding affinity for integrin α_V_β_3_ that act as vectors for delivery of chemotherapeutics [29]. These cyclic peptides have been used to ameliorate the cytotoxicity of drugs and reduce the side effects caused by toxicity without killing healthy cells. For instance, cyclic peptide conjugated antitumor small molecules, doxorubicin and paclitaxel, showed improved inhibition of growth and metastasis in cancer models [23].

Our results provide new information that the combination of sunitinib and integrin-targeted cyclic peptides exerts a synergistic anti-cancer effect for the treatment of EMT-mediated diseases, including pulmonary fibrosis and lung cancer metastasis.

## Materials and Methods

### Materials

Sunitinib was obtained from Selleck Chemicals (USA), and cRGDfK peptide was synthesized by Dr. Park (CHA Meditech Co., Ltd, Korea). Recombinant human TGF-β1 was purchased from R&D Systems, Inc. (USA). Fetal bovine serum (FBS), Dulbecco’s Modified Eagle’s Medium (DMEM), and antibiotics (100 U/mL penicillin and 100 μg/mL streptomycin) were purchased from Corning, Inc. (USA). Antibodies against p-ERK1/2, ERK1/2, p-Smad2/3, and Smad2/3 were purchased from Cell Signaling Technology, Inc. (USA). Antibodies against E-cadherin, N-cadherin, vimentin, TNIK, β-catenin, lamin B1, horseradish peroxidase (HRP)-conjugated secondary antibodies, and HRP-conjugated actin were purchased from Santa Cruz Biotechnology, Inc. (USA), and α-smooth muscle actin (α-SMA) was purchased from Abcam (UK).

### Cell Culture and Tissue Sample Preparation

Human NSCLC A549 cells were purchased from ATCC (USA) and H358 (KCLB No. 25807), H1299 (KCLB No. 25803), and human lung fibroblast IMR90 cells from Korean Cell Line Bank (KCLB No. 10186). A549, H358, and H1299 cells were maintained in DMEM and IMR90 cells in Minimum Essential Medium (MEM; Corning, USA) containing 10% heat-inactivated fetal bovine serum (FBS) and 1% antibiotics (100 U/mL penicillin and 100 μg/mL streptomycin). All cells were maintained at 37°C in a humidified atmosphere of 5% CO_2_. The use of human NSCLC patient’s tissue samples for this study was approved by the International Review Board of Eulji University (EU 18-14). The NSCL tissues were originated from human lung adenocarcinoma (10), bronchoalveolar carcinoma (3), squamous cell carcinoma (2), and large cell carcinoma (2).

### Cell Viability Assay

To assess cell viability, cells (5 × 10^3^ cells/well) were seeded in a 96-well plate for 24 h. Cells were treated with or without TGF-β1 (5 ng/mL) and sunitinib or cRGDfK for 24–72 h in culture media containing 10% FBS. Cell viability was assessed using the Cell Counting Kit-8 (Dojindo Molecular Technologies, Japan) according to the manufacturer’s instructions. The absorbance was measured with a Multiscan™ FC microplate photometer (Thermo Fisher Scientific, USA). Cell viability is presented as the percentage of control (untreated cells). Experiments were performed in triplicate.

### Molecular Docking

Molecular docking was carried out using Discovery Studio 2017 R2 and the crystal structure of TNIK (PDB ID 5AX9) (http://www.rcsb.org). The protein was prepared using the Prepare Protein module of Discovery Studio 2017 R2 under CHARMm force field [30] and default conditions. The TNIK binding site sphere was defined as a volume using the applicable module in Discovery Studio 2017 R2. The backbone carbonyl group of Glu106, the backbone nitrogen, and the carbonyl groups of Cys108 were selected as hydrogen bond constraints.

### RNA interference

A549 cells were transfected with TNIK siRNA or non-targeting control siRNA using the siRNA Reagent System (Santa Cruz Biotechnology, Inc., USA). After 48 h of incubation, the mRNA expression of Wnt target genes and mesenchymal marker genes by qRT-PCR analysis (see the “qRT-PCR” section below).

### TCF/LEF Reporter Assay

A549 cells were seeded in a 96-well plate for 24 h. Cells were treated with or without TGF-β1 (5 ng/mL) and sunitinib or cRGDfK. The relative TCF/LEF reporter activity was measured by the Cignal™ TCF/LEF Reporter Assay Kit (Qiagen, Germany) according to the manufacturer’s instructions. After 48 h, cells were washed with PBS, and then lysed using passive lysis buffer (Promega, USA). Luciferase activity was evaluated using the Dual Luciferase Reporter Assay kit (Promega, USA) and normalized by the total lysate of negative control-transfected cells. All experiments were performed in triplicate.

### Western blot Analysis

Serum-deprived A549 cells were treated with TGF-β1 (5 ng/mL) and sunitinib or cRGDfK for 72 h. Cytoplasmic and nuclear fractions were prepared from lysates using Nuclear and Cytoplasmic Extraction Reagent (Thermo Fisher Scientific, USA). Cytoplasmic or nuclear fractions were loaded on 8–15% polyacrylamide gels and transferred to nitrocellulose membranes. Specific primary antibodies were used to detect the expression of proteins. After incubation with HRP-conjugated secondary antibodies, the signals were visualized using Luminata™ Forte Western HRP Substrate (Merck Millipore, Germany). The band intensities were measured to determine the relative protein expression using X-ray films and development solution (Fujifilm, Japan). Actin or lamin B1 were used as loading controls. The detected bands were quantified based on the ImageJ software, and the relative ratio between each sample and loading controls was presented in the figures. The average band intensities of the independent three western blot results are shown in S1 Fig.

### Quantitative Real-Time PCR (qRT-PCR) Analysis

Cells were incubated in 60 mm^2^ dishes for 24 h. Total RNA was isolated using the AccuPrep® RNA Extraction Kit (Bioneer Corp., Korea) and cDNA synthesized from 1 μg of total RNA using oligo (dT) primers (Bioneer Corp., Korea) and the RocketScript™ Reverse Transcriptase Kit (Bioneer Corp., Korea). qRT-PCR was performed with the ExcelTaq 2X Q-PCR Master Mix (SMOBiO, Taiwan) and CFX96 ™ Real-Time System (Bio-Rad, USA). The cycling conditions were as follows: 95°C for 3 min followed by 40 cycles at 95°C for 15 s, 60°C for 30 s, and 72°C for 30 s. The primer sequences used in this study are provided in S1 Table. All reactions were run in triplicate, and data were analyzed using the 2^−ΔΔC^_T_ method [31]. The internal standard was GAPDH. Significance was determined by the Student’s *t*-test with GAPDH-normalized 2^−ΔΔC^_T_ values [31].

### Analysis of Combined Drug Effects

We analyzed the effects of the drug combination using the CalcuSyn software program (Biosoft, UK). To determine whether the result of treatment with the two compounds was additive or synergistic, we applied combination index (CI) methods derived from the median effect principle of Chou and Talalay [32]. The CI was calculated by the formula published by Zhao et al. [33]. A CI of 1 indicated an additive effect between the two compounds, a CI > 1 indicated antagonism, and a CI < 1 indicated synergism.

### Statistical Analysis

Data are presented as the mean ± SD of at least three independent experiments. Significant differences were evaluated by the One-way ANOVA with post-hoc Tukey HSD Test, with *P* < 0.05 considered significant. ^#^ *p* < 0.01 versus control, * *p* < 0.05, ** *p* < 0.01 versus treatment with TGF-β1 only.

## Results

### Sunitinib has high affinity for the ATP binding site of TNIK

We performed molecular docking studies to gain insight into the binding mode of sunitinib to TNIK at the atomic level. We confirmed that E106 and C108 in the hinge region interact with three hydrogen bonds (Fig 1A). The E106 and C108 backbone hydrogen bonds interact with the pyrrolidone of sunitinib at distances of 2.81 Å and 2.03 Å, respectively. The pyrrole of sunitinib interacts with C108 in the backbone at a distance of 2.91 Å. Sunitinib binding to TNIK is further stabilized by the CH/π interaction with the hydrophobic properties of TNIK A52, V31, L160, and V170.

**Figure 1.**
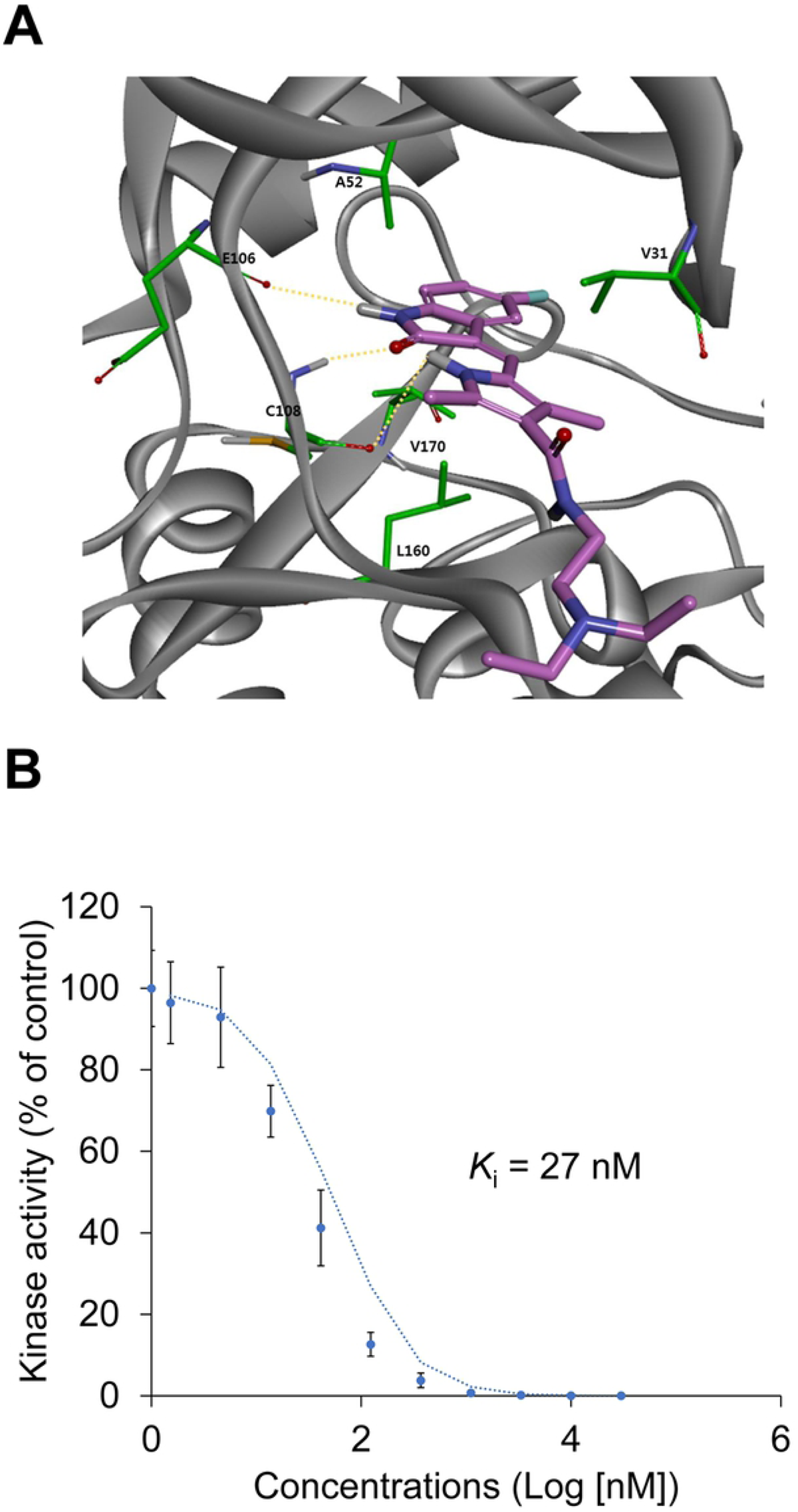
Binding mode and *K*_i_ of sunitinib for TNIK. (A) The proposed binding mode of sunitinib (pink) in the ATP-binding pocket of TNIK. TNIK and its interacting residues are represented by ribbons and sticks, respectively. Hydrogen bonding interactions appear as dashed yellow lines. (B) The binding constant, *K*_i_, of sunitinib for TNIK was determined using an ATP competition assay.

The inhibitory binding constant, *K*_i_, of sunitinib for TNIK was also calculated from triplicate, 11-point dose-response curves (Fig 1B). Sunitinib inhibited the binding of ATP to TNIK in a dose-dependent manner (*K*_i_ = 27 nM). These results indicate that the high affinity of sunitinib for TNIK inhibits ATP binding and leads to a decrease in TNIK kinase activity.

### Sunitinib inhibits TNIK-expressing NSCLC cell proliferation and TGF-β1-induced activation of Wnt signaling

To evaluate the inhibitory effect of sunitinib on TNIK-expressing NSCLC cell proliferation, we confirmed the mRNA expression of TNIK in NSCLC patient’s tissue cells and protein expression in normal lung fibroblast IMR90 and NSCLC A549, H1299, and H358 cells. TNIK mRNA expression was higher in NSCLC tissue cells than in normal lung tissues (Fig 2A). In addition, IMR90 cells did not express TNIK, but A549, H1299, and H358 cells did express the protein (Fig 2B). Sunitinib time- and dose-dependently inhibited the survival of A549, H1299, and H358 cells (Fig 2C-E). However, IMR90 cells were not significantly affected by sunitinib compared to NSCLC cells (Fig 2F). This result suggests that sunitinib has stronger anti-proliferative effect in TNIK-expressing NSCLC cells than normal lung fibroblasts.

**Figure 2.**
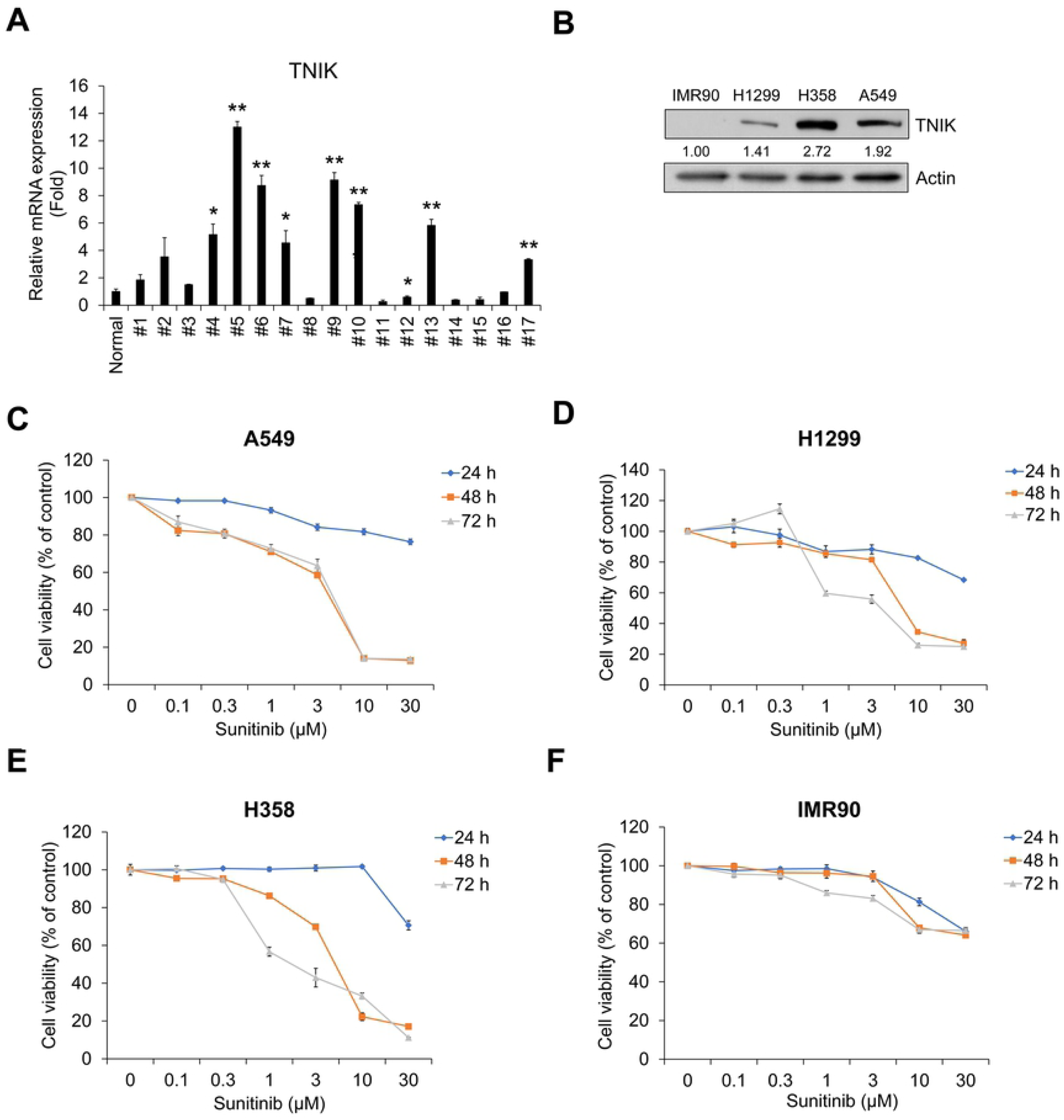
Sunitinib inhibits the proliferation of TNIK-expressing human NSCLC cells. (A) TNIK mRNA expression in NSCLC tissue cells from lung cancer patients. TNIK mRNA expression levels were measured by qRT-PCR analysis in pool normal lung tissue cells (n = 8) and in NSCLC cells from patients with lung cancer (n = 17). The normal group represents the average qRT-PCR results obtained from eight healthy lung tissue cells without cancer cells. Data are represented as mean ± SD. Experiments were performed in triplicate. *p < 0.05 and **p < 0.001 (vs. control). (B) The protein expression of TNIK in normal fibroblast IMR90 and NSCLC H1299, H358, and A549 cells was measured by Western blot. Actin was used as a loading control. (C–F) A549 (C), H1299 (D), and H358 (E) cells and IMR90 (F) cells were treated with sunitinib for 24–72 h. After incubation, cell viability was measured by CCK-8 assay. Experiments were performed in triplicate. Data represent mean ± SD.

Next, we confirmed the mRNA expression using qRT-PCR analysis whether silencing of TNIK affect the transcriptional activity of TNIK and Wnt target genes in A549 cells. Silencing of TNIK inhibited TGF-β1-induced mRNA expression of *TNIK, CTNNB1, TCF4*, and *c-MYC* (S2 Fig). Based on these results, we assessed the effect of sunitinib on the TGF-β1-induced activation of Wnt signaling pathways and TNIK expression. We previously demonstrated that TGF-β1 activates TCF4-mediated transcription and TNIK expression [18]. To confirm the effect of sunitinib on TCF4-mediated Wnt signaling pathways, we assessed the TGF-β1-induced β-catenin luciferase activity and transcriptional activity of Wnt target genes. TGF-β1-induced β-catenin luciferase activity was inhibited by sunitinib in a dose-dependent manner (Fig 3A). Activation of TNIK is mediated by the binding of β-catenin to TCF4 and subsequent phosphorylation of TCF4, which activates Wnt target gene expression [16,17]. We confirmed that TGF-β1 induced the transcription of TNIK and Wnt target genes. Increased mRNA expression of TNIK and Wnt target genes was strongly inhibited by sunitinib (Fig 3B). Sunitinib also suppressed cytosolic β-catenin and nuclear TNIK and β-catenin protein expression (Fig 3C). These results indicate that sunitinib has an anti-proliferative effect and inhibits TGF-β1-induced activation of Wnt signaling in NSCLC A549 cells.

**Figure 3.**
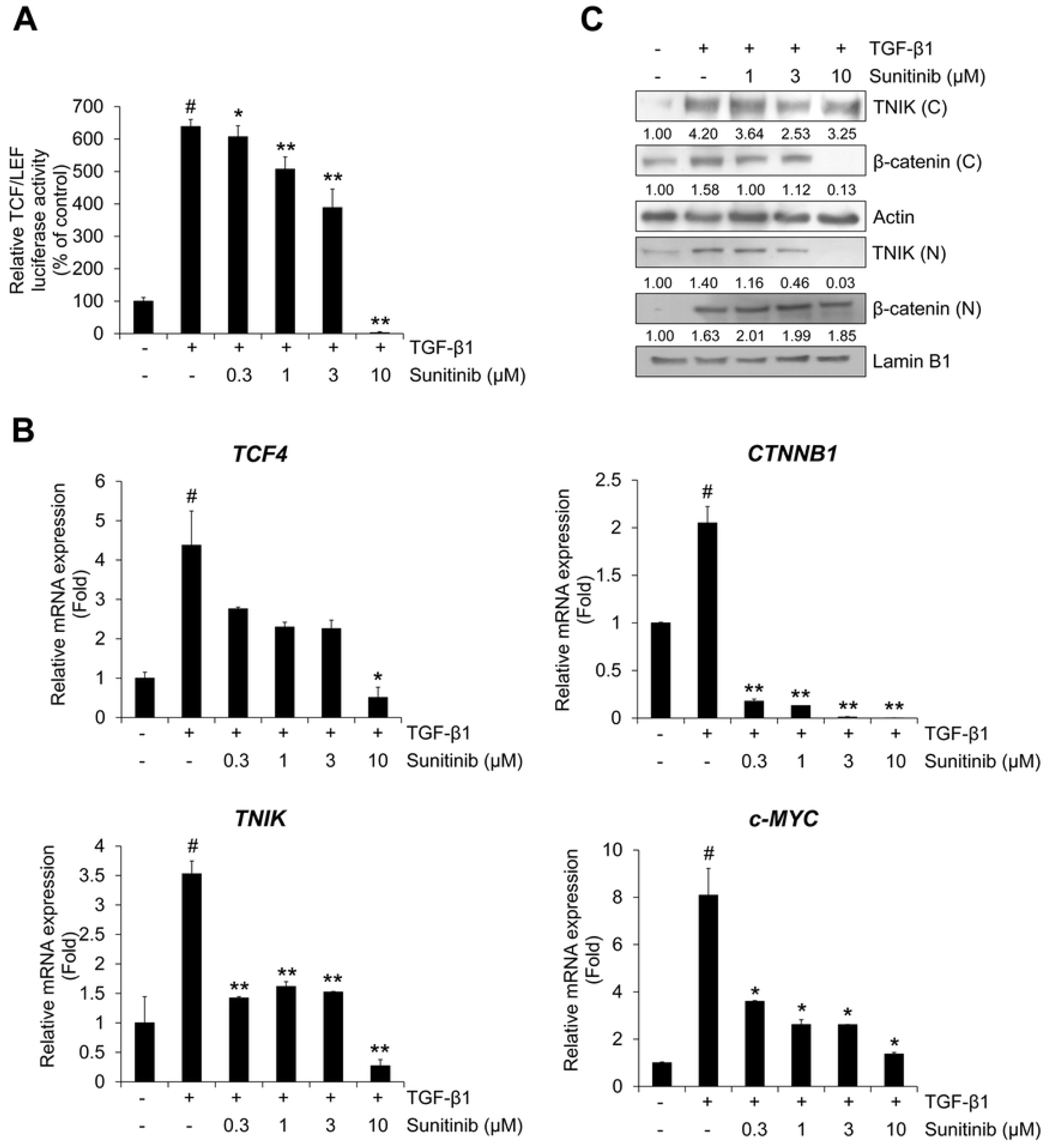
Sunitinib inhibits TGF-β1-induced Wnt signaling. (A) Effect of sunitinib on TCF4 transcriptional activity. A549 cells were treated with TGF-β1 (5 ng/mL) or its combination with sunitinib for 48 h before measuring TOPflash luciferase activity. FOPflash-normalized TOPflash luciferase activity is represented as the relative TCF/LEF luciferase activity. (B) A549 cells were pretreated with sunitinib for 3 h, and then incubated with TGF-β1 (5 ng/mL) for 1 h. After RNA extraction and cDNA synthesis, the mRNA expression of TNIK and Wnt target genes was measured by qRT-PCR. (C) The protein levels of TNIK and β-catenin in cytosolic and nuclear fractions were measured by Western blot. Actin and lamin B1 were used as loading controls for the cytosolic and nuclear fractions, respectively. The results are representative of triplicate experiments. ^#^ *p* < 0.01 versus control; * *p* < 0.05, ** *p* < 0.01 versus treatment with TGF-β1 only.

### Sunitinib inhibits TGF-β1-induced EMT and Smad/non-Smad signaling

As in the above result, we also confirmed the effect of TNIK silencing on EMT phenotype by qRT-PCR analysis. Silencing of endogenous TNIK suppressed TGF-β1-induced mRNA expression of mesenchymal maker genes *CDH2* and *VIM* encoding N-cadherin and vimentin, respectively (S3 Fig). However, transfection of TNIK siRNA did not significantly affect the mRNA expression of *CDH1* encoding E-cadherin. Based on these results, to verify the inhibitory effect of sunitinib on TGF-β1-induced EMT, we evaluated the protein expression of epithelial and mesenchymal markers. TGF-β1 suppressed the expression of epithelial marker E-cadherin, but stimulated mesenchymal markers N-cadherin, vimentin, and α-smooth muscle actin (SMA) [1,2]. Sunitinib dose-dependently inhibited TGF-β1-induced protein expression of EMT markers in A549 cells (Fig 4A). We also confirmed these inhibitory effects of sunitinib on the TGF-β1-mediated transcription of mesenchymal marker genes *CDH2* and *VIM*, encoding N-cadherin and vimentin, respectively (Figs 4B and 4C); the TGF-β1-induced mRNA expression of both *CDH2* and *VIM* was suppressed.

**Figure 4.**
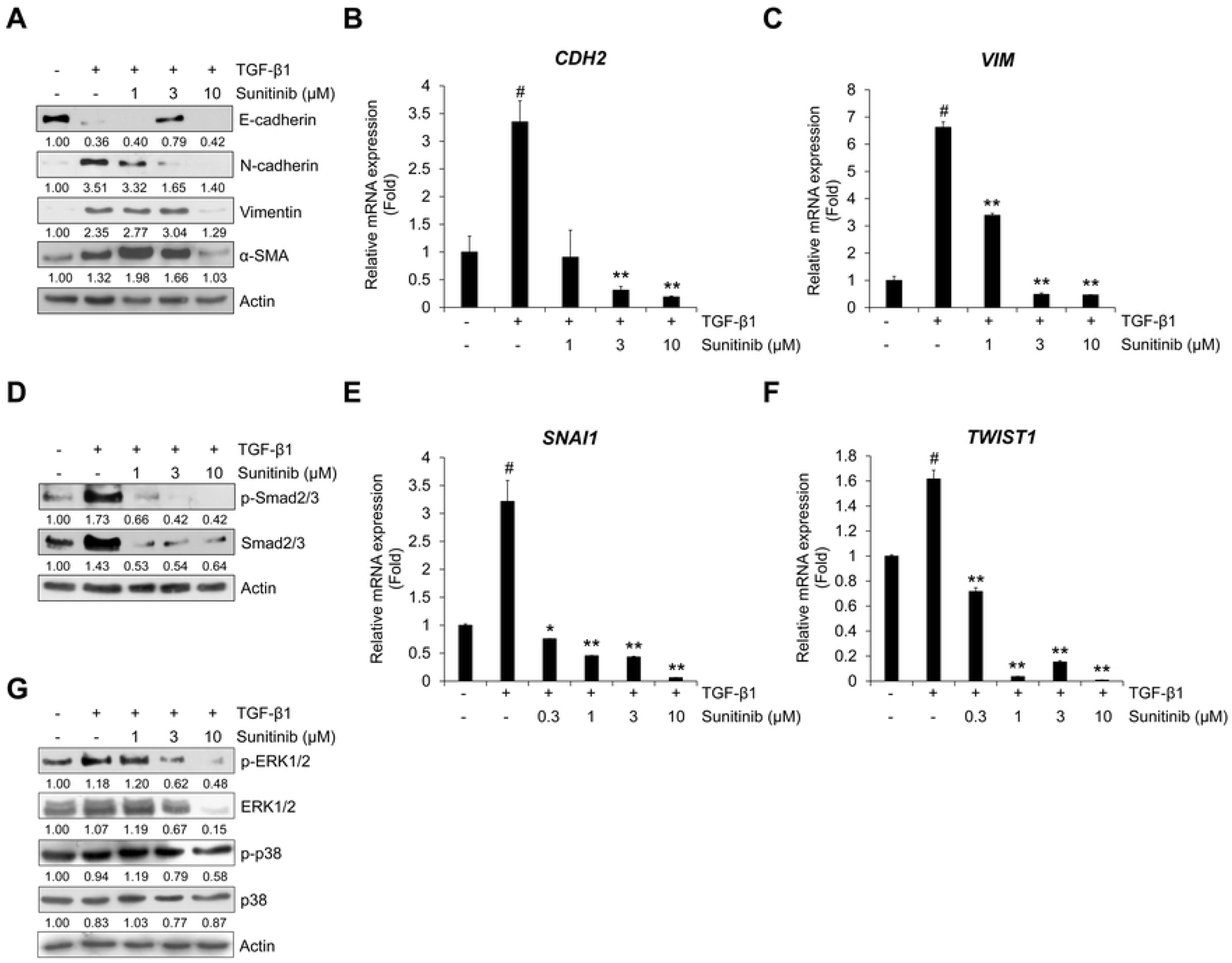
Sunitinib inhibits TGF-β1-induced expression of EMT markers and Smad signaling. (A) Serum-deprived A549 cells were treated with TGF-β1 (5 ng/mL) or its combination with sunitinib for 72 h. The protein expression of TGF-β1-mediatead EMT markers was evaluated by Western blot. Actin was used as a loading control. (B and C) Serum-deprived A549 cells were pretreated with sunitinib for 3 h, and then incubated with TGF-β1 (5 ng/mL) for 48 h. After RNA extraction and cDNA synthesis, we performed qRT-PCR to measure mRNA expression of *CDH2* and *VIM* using GAPDH as an internal control. (D) Effects of sunitinib on TGF-β1-activated Smad signaling. A549 cells were treated with TGF-β1 (5 ng/mL) or its combination with sunitinib for 24 h, and then the levels of p-Smad2/3 and endogenous Smad2/3 in cytosolic fraction were evaluated by Western blot. (E and F) The transcriptional activity of *SNAI1* and *TWIST1* was measured by qRT-PCR using GAPDH as an internal control. Serum-deprived A549 cells were pretreated with sunitinib for 3 h, and then incubated with TGF-β1 (5 ng/mL) for 1 h. (G) The effect of sunitinib on TGF-β1-induced phosphorylation of ERK1/2 and p38 was determined by Western blot. Actin was used as a loading control. ^#^ *p* < 0.01 versus control, * *p* < 0.05, ** *p* < 0.01 versus treatment with TGF-β1 only.

To elucidate the inhibitory effect of sunitinib on the TGF-β1-mediated EMT-related signaling pathway, we examined the level of TGF-β1-induced phosphorylation of Smad2/3 involved in the EMT process and the transcriptional activity of Smad signaling target genes *SNAI1* (also referred to as *Snail*) and *TWIST1* (also referred to as *Twist*), upregulating mesenchymal markers and downregulating epithelial markers. Sunitinib inhibited TGF-β1-induced phosphorylation of Smad2/3 (Fig 4D). We also confirmed that TGF-β1-induced mRNA expression of *SNAI1* and *TWIST1* was suppressed by sunitinib (Figs 4E and 4F). These results demonstrate that the TGF-β1-induced EMT phenotype was strongly attenuated by sunitinib through suppression of Smad-mediated signaling.

TGF-β can also regulate EMT and invasion through non-Smad signaling pathways, including MAP kinase, Ras-ERK, and JNK [34-36]. In a previous study, we investigated the induction of ERK1/2 phosphorylation by TGF-β1 treatment [18]. We evaluated the inhibitory effects of sunitinib on TGF-β1-induced activation of ERK. As shown in Fig 4G, sunitinib inhibited the TGF-β1-induced phosphorylation of ERK1/2, suggesting that sunitinib can suppress the TGF-β-mediated EMT process through the inhibition of non-Smad signaling, as well as the Smad-signaling pathway.

### cRGDfK inhibits TGF-β1-induced expression of mesenchymal markers in A549 cells

To improve the effect of sunitinib, A549 cells were co-treated with sunitinib and cyclic-RGDfK peptide (cRGDfK) (Fig 5A and S4 Fig). First, to confirm the therapeutic potency of cRGDfK as an integrin α_V_β_3_ antagonist in NSCLC cells, we checked the mRNA expression of *ITGAV* and *ITGB3* encoding integrin α_V_ and integrin β_3_, respectively. As shown in Fig 5B, A549 cells showed more *ITGAV* and *ITGB3* mRNA expression than those of IMR90, H1299 and H358 cells. Next, to assess the cytotoxicity of cRGDfK, we conducted cell viability assays in both NSCLC cell lines and normal fibroblast IMR90 cells. cRGDfK suppressed proliferation of NSCLC cells, but IMR90 cells were not significantly affected (Figs 5C–5F). In addition, cRGDfK had an inhibitory effect on TGF-β1-induced expression of TNIK and β-catenin in A549 cells (S5 Fig).

**Figure 5.**
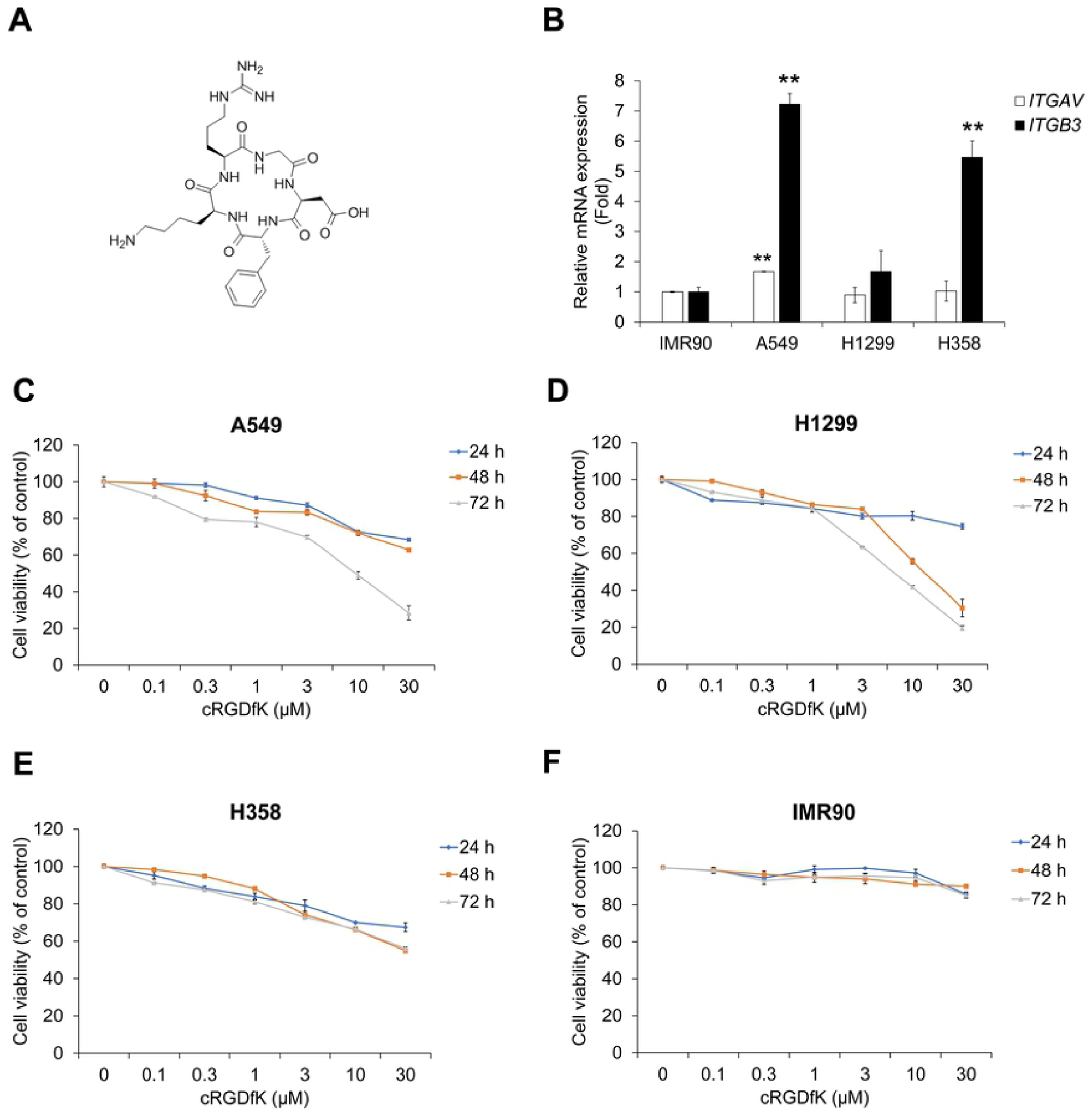
Treatment with cRGDfK inhibits proliferation of human NSCLC cells. (A) Chemical structure of cRGDfK. (B) The relative mRNA expression of integrin α_V_ and β_3_ in human NSCLC cells. (C–F) NSCLC A549 (C), H1299 (D), and H358 (E) cells and normal fibroblast IMR90 cells (F) were treated with cRGDfK for 24–72 h. After incubation, cell viability was measured by CCK-8 assay. Experiments were performed in triplicate. Data represent mean ± SD.

We assessed whether cRGDfK inhibits the expression of TGF-β1-induced EMT markers in A549 cells. As shown in Fig 6A, the TGF-β1-induced reduction of E-cadherin expression was not restored by cRGDfK, but the increase in N-cadherin, vimentin and α-SMA was ihibited. This result was confirmed by the mRNA expression of *CDH2* and *VIM* encoding N-cadherin and vimentin, respectively. However, there were no concentration-dependent decreases in mRNA expression with cRGDfK treatment, and cRGDfK inhibited the TGF-β1-induced transcriptional activity of *CDH2* and *VIM* (Figs 6B and 6C). The protein expression of phosphorylated Smad2/3 in the cytosol was also inhibited by cRGDfK (S6 Fig). Treatment with cRGDfK strongly suppressed TGF-β1-induced transcriptional activity of *SNAI1* and *TWIST1* (Figs 6D and 6E). These results demonstrate that cRGDfK has an inhibitory effect on the TGF-β1-mediated EMT process in NSCLC A549 cells.

**Figure 6.**
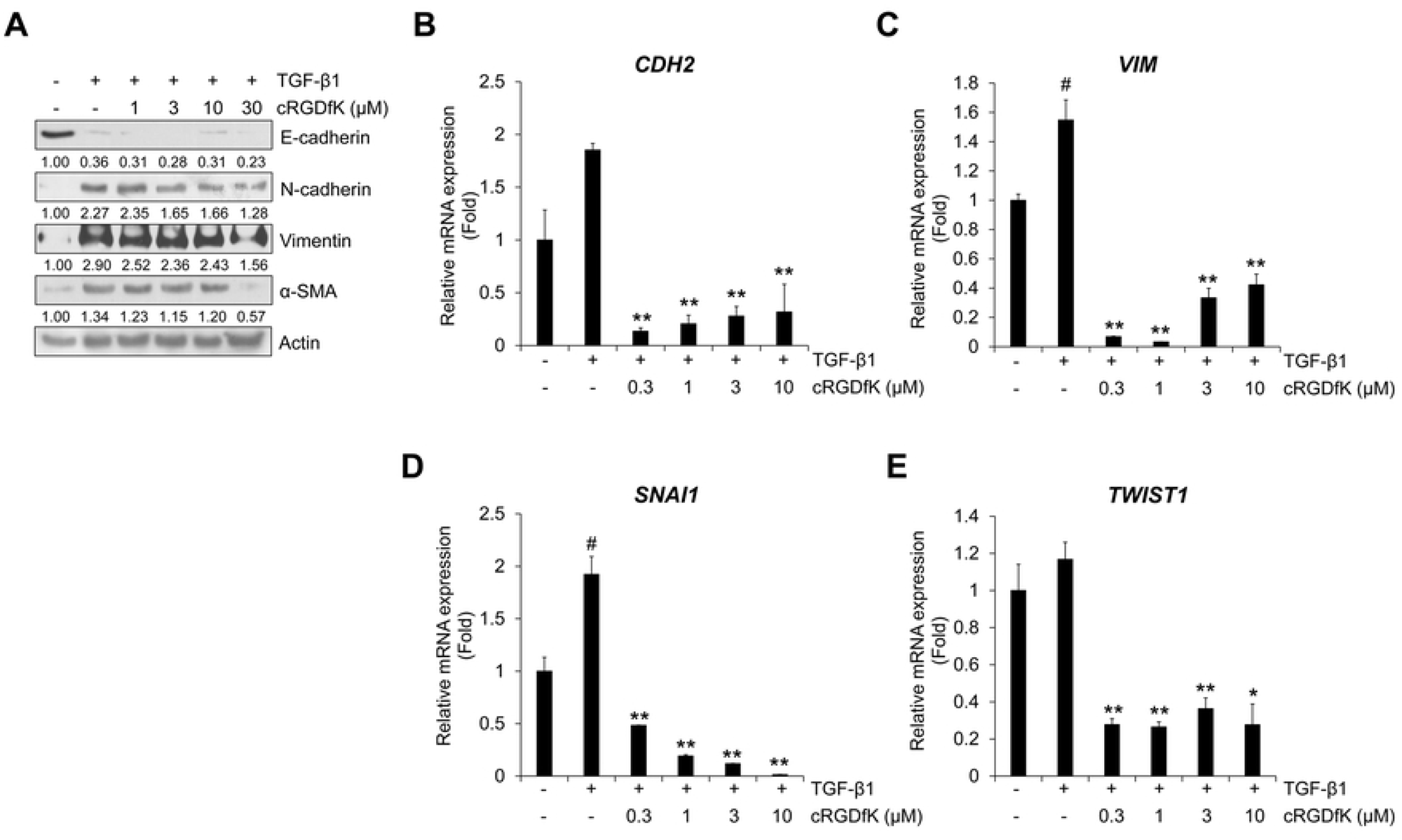
Treatment with cRGDfK inhibits TGF-β1-induced expression of EMT markers. (A) Serum-deprived A549 cells were treated with TGF-β1 (5 ng/mL) or its combination with cRGDfK for 72 h. The protein expression of TGF-β1-mediatead EMT markers was detected by Western blot. Actin was used as a loading control. (B–E) Serum-deprived A549 cells were pretreated with cRGDfK for 3 h, and then treated with TGF-β1 (5 ng/mL) for 48 h (B and C) or 1 h (D and E). The mRNA expression of *CDH2* and *Vim* (B and C) or *Snail* and *Twist* (D and E) was measured by qRT-PCR using GAPDH as an internal control. Experiments were performed in triplicate. Data represent mean ± SD. ^#^ *p* < 0.01 versus control, * *p* < 0.05, ** *p* < 0.01 versus treatment with TGF-β1 only.

### Combined treatment with cRGDfK improves the inhibitory effect of sunitinib on cell viability and TGF-β1-induced EMT process in A549 cells

To evaluate the effect of cRGDfK on the TGF-β1-induced EMT process in NSCLC A549 cells with sunitinib, we evaluated cell viability. Compared to the single treatment with sunitinib, combined treatment with cRGDfK increased the inhibitory effect on A549 cell proliferation (Fig 7A). This result was confirmed by calculation of the combination index (CI) using the raw data (Fig 7A), which evaluated a synergistic effect at concentrations > 0.3 μM (Table 1). In addition, we confirmed the combination effect on the TGF-β1-induced protein expression of EMT markers and mRNA expression of target genes. Reduced expression of E-cadherin and *CDH1* induced by TGF-β1 was recovered by combined treatment with sunitinib and cRGDfK (Figs 7B and 7C). The protein expression of N-cadherin and vimentin and transcriptional activity of *CDH2* were synergistically inhibited by combined treatment (Figs 7B, 7D and 7E). The inhibitory effect of sunitinib on the mRNA expression of Smad signaling target genes *SNAI1* and *Twist* was also enhanced by cRGDfK (Figs 7F and 7G).

**Table 1.**
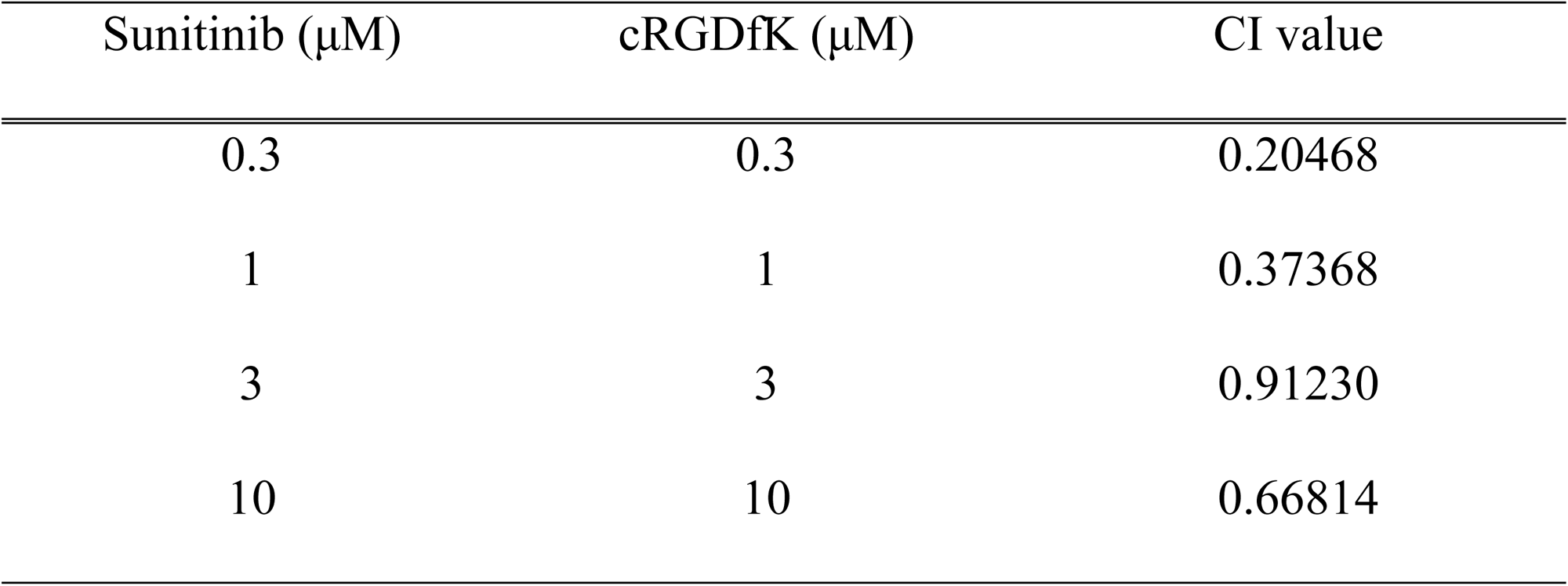
Combination index (CI) values for the two-drug combination against A549 cell viability.

**Figure 7.**
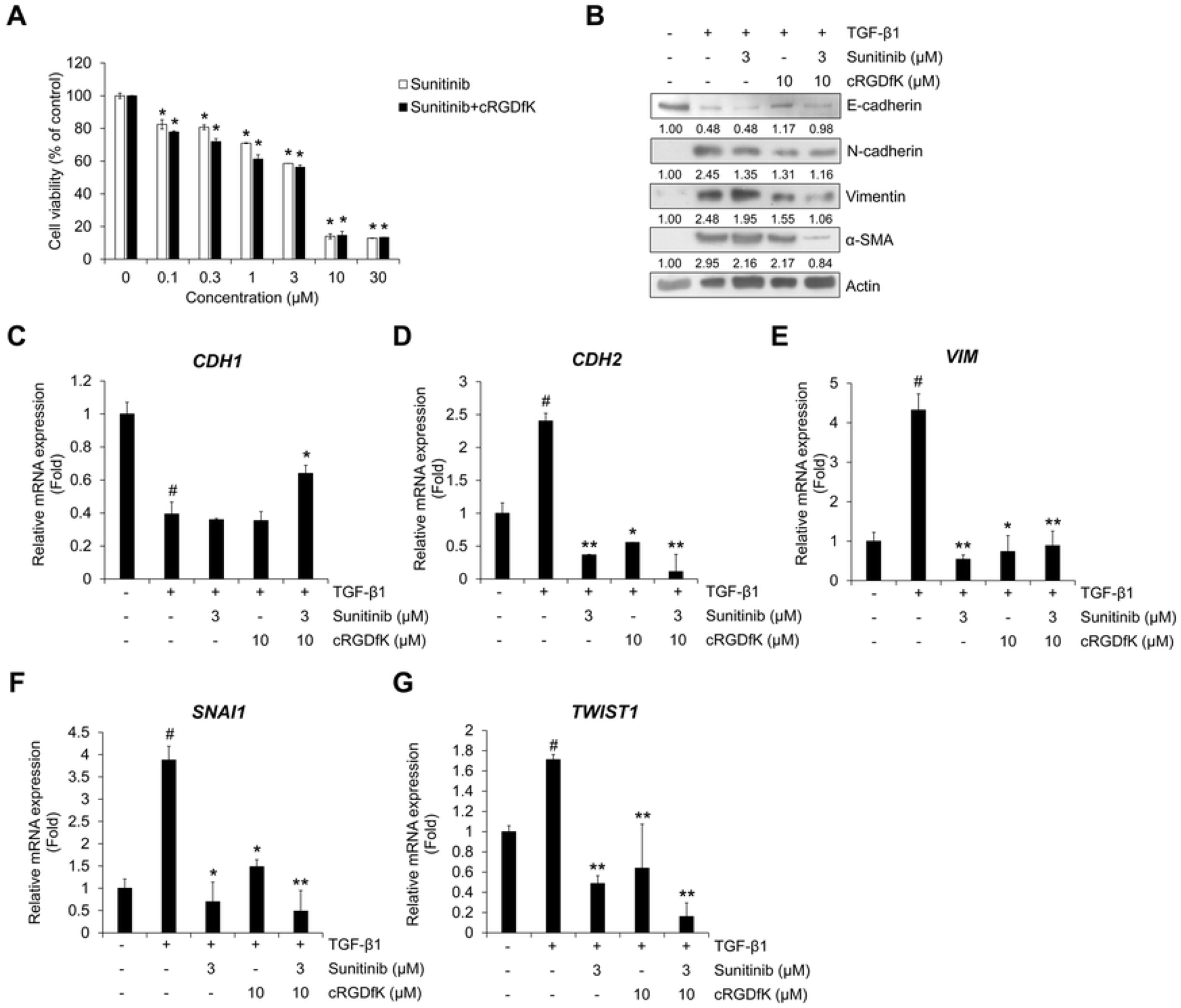
Combination of sunitinib with cRGDfK shows enhanced inhibitory effects on cell viability and TGF-β1-induced EMT expression. (A) A549 cells were treated with sunitinib or a combination of sunitinib plus cRGDfK for 24 h before measuring cell viability. Data represent mean ± SD. *p < 0.01 vs. sunitinib-free controls. (B) Serum-deprived A549 cells were pretreated for 3 h with sunitinib (3 μM) and cRGDfK (10 μM) individually or in combination, and then incubated with TGF-β1 (5 ng/mL) for 72 h. The expression of EMT markers was measured by Western blot. Actin was used as a loading control. (C–G) A549 cells were pretreated with sunitinib or a combination of sunitinib plus cRGDfK for 3 h and incubated with TGF-β1 (5 ng/mL) for 48 h (C–E) or 1 h (F and G). The mRNA expression of the indicated genes was measured by qRT-PCR using GAPDH as an internal control. Experiments were performed in triplicate. Data represent mean ± SD. ^#^ *p* < 0.01 versus control, * *p* < 0.05, ** *p* < 0.01 versus treatment with TGF-β1 only.

These results suggest that cRGDfK exerts an enhancing effect on A549 cell viability and TGF-β1-induced EMT as a targeting peptide that increases the drug efficacy of sunitinib in lung cancer cell death and the EMT process.

## Discussion

Loss of epithelial characteristics and acquisition of mesenchymal features in the EMT is a key mechanism for metastatic and invasive changes in cancer cells [1,2]. EMT also stimulates anti-apoptotic signals that confer resistance to chemotherapy in NSCLC cells [36]. Thus, inhibition of EMT during tumor progression is a very important strategy for the treatment and prevention of NSCLC, together with the induction of cancer cell death. Here, we showed that TGF-β1 strongly induces the EMT and Smad/non-Smad signaling pathways of NSCLC A549 cells, and as a receptor tyrosine kinase (RTK) inhibitor sunitinib significantly inhibited these inductions. We previously demonstrated that TNIK is involved in TGF-β1-induced EMT, migration, and invasion of NSCLC A549 cells [18]. In this study, inhibition of TNIK by siRNA or a specific TNIK inhibitor, KY-05009, suppressed cell proliferation and the TGF-β1-induced EMT process. As sunitinib also has been shown to inhibit TNIK kinase activity [21], and to address the relevance between TNIK inhibition and TGF-β1-induced EMT, we evaluated the effect of sunitinib as an anti-cancer agent inhibiting TNIK kinase activity and the EMT in NSCLC cells. As expected, sunitinib was found to have three hydrogen bonds with the hinge region of TNIK, which allows inhibition of kinase activity by strong interactions with TNIK. Therefore, we hypothesized that inhibition of TNIK by sunitinib can regulate TGF-β1-induced EMT and invasion in NSCLC cells. The results show that sunitinib has the potential to inhibit NSCLC cell proliferation and TGF-β1-induced EMT through Smad/non-Smad signaling and TNIK-mediated Wnt signaling.

Although several types of RTK inhibitors, such as erlotinib and gefitinib, are used to treat NSCLC, the EMT process in NSCLC is a major cause of resistance to long-term chemotherapy [36]. Therefore, to overcome the limitation of single drug chemotherapy, recent trends in NSCLC therapy have sought combinations with other NSCLC treatments that may increase drug efficacy, including immunotherapy, radiotherapy, and targeted therapy [37]. The use of peptides in these combination therapies is a good way to reduce side effects that can occur with multiple drug use. In particular, cyclic peptides have been actively studied recently as biochemical tools and therapeutic agents because of their excellent stability, high resistance to exogenous peptidases, and high affinity and selectivity for binding to target biomolecules [38]. Cyclic pentapeptides containing the RGD motif (cRGDfK) are used to improve the drug delivery and target selectivity of anti-cancer drugs [34,39,40]. Many studies have demonstrated the potential of RGD-containing peptides as vectors for delivering anticancer agents. In particular, cyclic peptides such as cRGDfK and cRGDyK showed high binding affinity and selectivity for intergin α_V_β_3_. Thus, the use of cyclic peptides has been as one of the methods of delivery of therapeutic small molecules such as doxorubicin and paclitaxel in cancers [23]. However, the combined effect of RTK inhibitors and cRGDfK in the EMT process in the tumor microenvironment has not been fully investigated. Based on previous studies, we investigated whether cRGDfK enhances the inhibitory effect of sunitinib on cell proliferation and TGF-β1-induced EMT in NSCLC cells. We demonstrated that sunitinib itself exerts inhibitory effects on TNIK kinase activity and the TGF-β1-induced EMT in A549 cells. However, the inhibitory effects of sunitinib were increased by co-treatment with cRGDfK. A549 cell proliferation and the mRNA and protein expression of EMT markers were synergistically inhibited by sunitinib and cRGDfK co-treatment. These results demonstrate that the effect of cRGDfK, as a targeting peptide, can enhance the permeability of sunitinib to NSCLC cells in anti-EMT or anti-migratory/invasive therapy.

Recent reports suggest new methods to improve the therapeutic efficacy of tyrosine kinase inhibitors (TKIs) for NSCLC using combination strategies [37,41]. Two or more TKIs or TKIs and other target therapies (e.g., immunotherapeutics) have been used in the combination regimens, but there is not much known to be used in combination with peptides. In addition, the use of TNIK inhibitors and cyclic peptides for the EMT process in NSCLC progression has not yet been studied. In this regard, the present study suggests that a combination of cRGDfK with TKIs may be a new applicable method for the treatment of NSCLC. In particular, the development of new drugs in the modern biopharmaceutical industry is increasingly challenged by the time and expense of R&D expenditures. To reduce the R&D costs, companies are diversifying how they develop new drugs. Drug repositioning is a new method to search existing drugs for the application of new indications [42,43]. The advantage of drug repositioning is that it reduces the time and cost of developing new drugs, in that safety and toxicity assessments have already been completed. Thus, the use of TKIs and cRGDfK in NSCLC could increase the therapeutic efficacy without the development of new therapies by using existing therapeutic agents and the synergistic effect of drug efficacy.

Although studies on the improvement of anticancer drug efficacy using cyclic peptides have not been actively conducted, our results suggest that cyclic peptides may help prevent the progression of NSCLC using TKIs as therapeutic agents. We plan to conduct studies on the anticancer effect using cyclic peptides in many cancers other than NSCLC.

## Acknowledgments

The biospecimens and data used for this study were provide by the Biobank of Chungnam National University Hospital, a member of the Korea Biobank Network.

## Supporting information

**S1 Fig. The average band intensities of the independent three western blot results in Fig. 2B, 3C, 4A, 4D, 4G and 6A**. The detected bands were quantified based on the ImageJ software, and the relative ratio between each sample and loading controls was presented in the figures.

**S2 Fig. The mRNA expression of TNIK and Wnt target genes in A549 cells**. A549 cells were transfected with non-targeting control siRNA or TNIK siRNA. Transfected cells were treated with TGF-β1 (5 ng/mL) for 48 h. The mRNA expression was meausred by qRT-PCR analysis. Experiments were performed in triplicate. Data represent mean ± SD.

**S3 Fig. The mRNA expression of EMT marker genes in A549 cells**. A549 cells were transfected with non-targeting control siRNA or TNIK siRNA. Transfected cells were treated with TGF-β1 (5 ng/mL) for 48 h. The mRNA expression was meausred by qRT-PCR analysis. Experiments were performed in triplicate. Data represent mean ± SD.

**S4 Fig. Synthetic scheme for cyclic pentapeptide, c(RGDfK)** (**4**). The protected linear pentapeptide (**1**) bound to the resin was synthesized using the Fmoc solid phase peptide synthesis (SPPS) method. The linear peptide (**2**) was cleaved from the resin without affecting other protecting groups by using acetic acid/TFE/CH_2_Cl_2_ (1:1:3 ratio) solution. Finally, cyclic pentapeptide c(RGDfK) (**4**) was obtained by head-to tail cyclization under T3P, TEA, DAMP and elimination of the protecting group.

**S5 Fig. Western blot analysis of cytosolic TNIK and β-catenin expression**. Serum-deprived A549 cells were treated with TGF-β1 (5 ng/mL) or its combination with cRGDfK for 72 h. Actin was used as a loading control.

**S6 Fig. Western blot analysis of the effect of cRGDfK on TGF-β1-induced Smad-and non-Smad signaling in A549 cells**. Serum-deprived A549 cells were treated with TGF-β1 (5 ng/mL) or its combination with cRGDfK for 48 h (p-Smad2/3) or 72 h (p-ERK1/2 and p-p38). Actin was used as a loading control.

**S1 Table. Primer sequences used in this study**.

